# BigStitcher: Reconstructing high-resolution image datasets of cleared and expanded samples

**DOI:** 10.1101/343954

**Authors:** David Hörl, Fabio Rojas Rusak, Friedrich Preusser, Paul Tillberg, Nadine Randel, Raghav K. Chhetri, Albert Cardona, Philipp J. Keller, Hartmann Harz, Heinrich Leonhardt, Mathias Treier, Stephan Preibisch

## Abstract

New methods for clearing and expansion of biological objects create large, transparent samples that can be rapidly imaged using light-sheet microscopy. Resulting image acquisitions are terabytes in size and consist of many large, unaligned image tiles that suffer from optical distortions. We developed the BigStitcher software that efficiently handles and reconstructs large multi-tile, multi-view acquisitions compensating all major optical effects, thereby making single-cell resolved whole-organ datasets amenable to biological studies.

---

Sample clearing [1, 2] and expansion microscopy (ExM) [3] are powerful protocols that create large, transparent volumes of whole tissues and organisms. Using light-sheet microscopy [4–6], these samples can be imaged with subcellular resolution in their entirety within a few hours [7]. These acquisitions have the potential to be powerful tools for whole-tissue and whole-organism studies since they preserve endogenous fluorescent proteins and are compatible with most staining methods (**Supplementary Fig. 1**).

However, raw data acquired by the microscope is not directly suitable for visualization and analysis. Many large, overlapping three-dimensional (3d) image tiles are collected that amount to many terabytes in size and require image alignment (**Fig. 1d-n**). Due to sample-induced scattering of the light-sheet in the direction of illumination [8], 3d image tiles are typically acquired twice while alternating illumination from opposing directions to achieve full coverage (**Fig. 1d** and **Supplementary Fig. 2**). Similarly, emitted light is distorted by the sample, effectively limiting maximal imaging depth at which useful data can be collected (**Fig. 1n**). Additionally, sample-induced light refractions cause depth- and wavelength-dependent aberrations in the acquired images (**Fig. 1j,k**). To reconstruct and make these complex datasets easily accessible we developed the BigStitcher software. It enables interactive visualization using BigDataViewer [9], fast and precise alignment, real-time fusion, deconvolution, as well as support for alignment of multi-tile acquisition taken from different physical orientations, so-called multi-tile *views*, thereby effectively doubling the size of specimens that can be imaged (**Fig. 1n**), while further orthogonal views can render the resolution isotropic.

**Figure 1:**
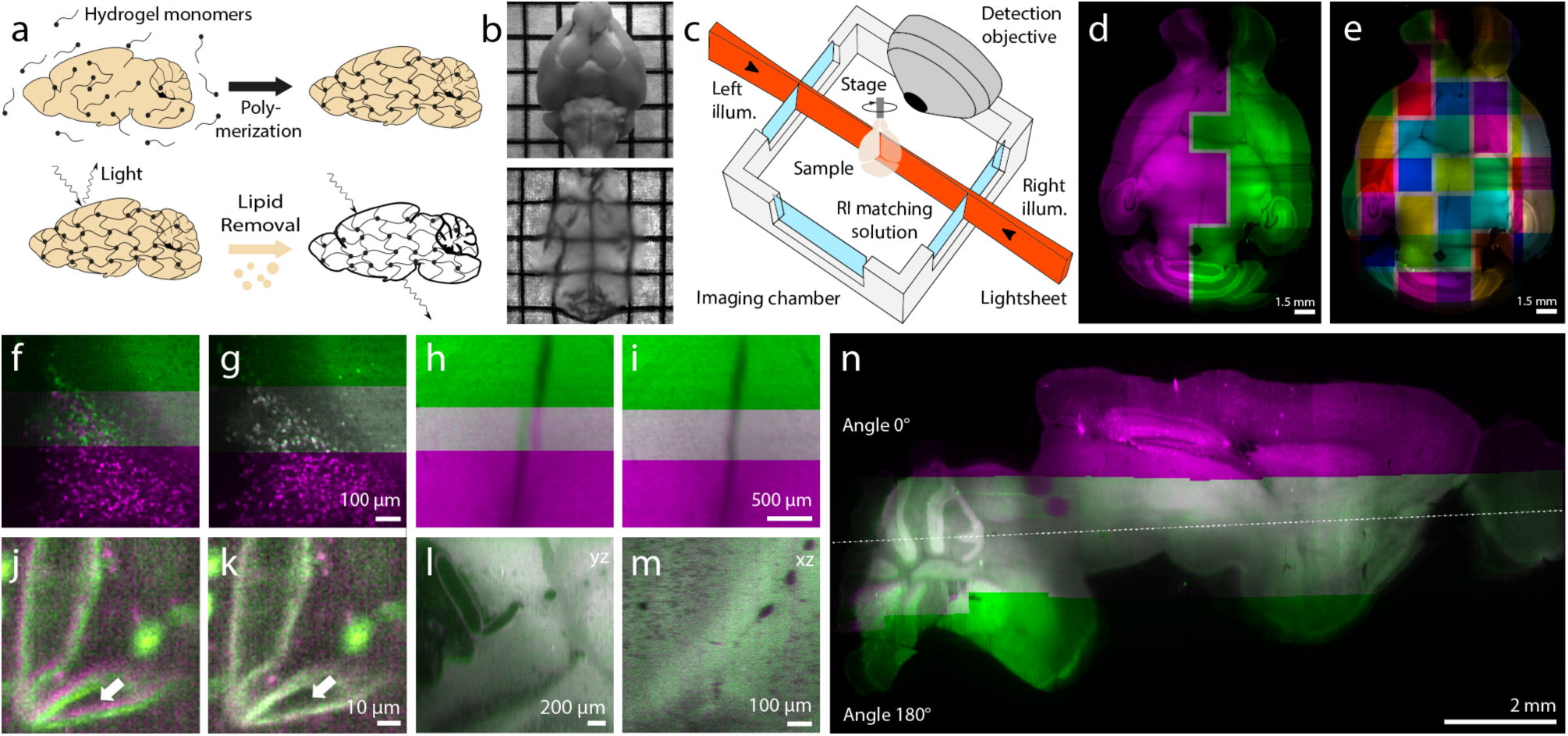
BigStitcher Principles. **(a)** schematic of the CLARITY sample clearing process. **(b)** adult mouse brain before (up) and after (down) clearing. **(c)** general layout of the type of light-sheet microscope used for most acquisitions [beads] **(d)** single slice through an entire adult mouse brain acquired with dual-sided illumination. Pink (left illum.) and green (right illum.) image tiles highlight the illumination direction that was automatically selected for each tile. **(e)** overview of an entire section of an acquired adult mouse brain, different colors highlight individual image tiles (each 1920×1920×770 pixels). **(f-i)** illustration of the result of the image stitching from a Bsx^H2BeGFP^ brain using phase correlation before (f,h) and after stitching (g,i). **(j,k)** the effect of ICP refinement on two different channels with sufficient autofluorescence visible in both channels. Arrows highlight significant difference before (j) and after (k) refinement. **(l,m)** quality of the multi-view reconstruction for two overlapping multi-tile views at 0 (magenta) and 180 (green) degrees for an axial vs. axial (l) and lateral vs. axial (m) view. **(n)** one slice through an entire adult mouse brain (2.24TB raw data), both views are shown in axial orientation looking along the rotation axis of the microscope. The dotted line illustrates the middle of the section.

Microscopy acquisitions are saved in a multitude of vendor-specific and custom formats. We developed an extendable, user-friendly interface that automatically imports almost any format and extracts relevant metadata such as illumination directions, sample rotation, and approximate image positions using Bioformats [10] (**Supplementary Note 1**). Alternatively, the importer supports interactive placement of image tiles using regular grids or simple text file-based definitions (**Supplementary Fig. 3**). BigStitcher accesses image data through memory-cached, virtual loading [11], optionally combined with virtual flatfield correction (**Supplementary Fig. 4** and **Supplementary Note 2**). Performance is optimal when images are stored using a multiresolution, blocked, compressed format enabling interactive visualization, processing and interaction with terabyte-sized image datasets. The importer therefore supports resaving single-block images into the BigDataViewer HDF5 format [9].

Although samples are highly transparent (**Fig. 1b**), light scattering is an issue when imaging centimeters deep into fixed tissue. Dual-sided light-sheet illumination (**Fig. 1c**) is able to significantly increase the sample size for which high resolution image data can be collected laterally by imaging each 3d image tile twice using left-sided and right-sided illumination (**Fig. 1d**). Since most tiles only hold useful information from either direction, we automatically suggest the best illumination direction for each tile by estimating image sharpness at the lowest pre-computed resolution level (**Fig. 1d** and **Supplementary Fig. 2**).

To compute locations for every image tile we developed a new image stitching algorithm. It is tailored for very large datasets and can deal with acquisitions arranged in non-regular grids (**Fig. 2a**) containing empty images and multiple independent samples (**Supplementary Fig. 5**). We therefore compute overlaps between all pairs of overlapping tiles, identify incorrect pairwise overlaps, and compute globally optimal positions for all tiles.

**Figure 2:**
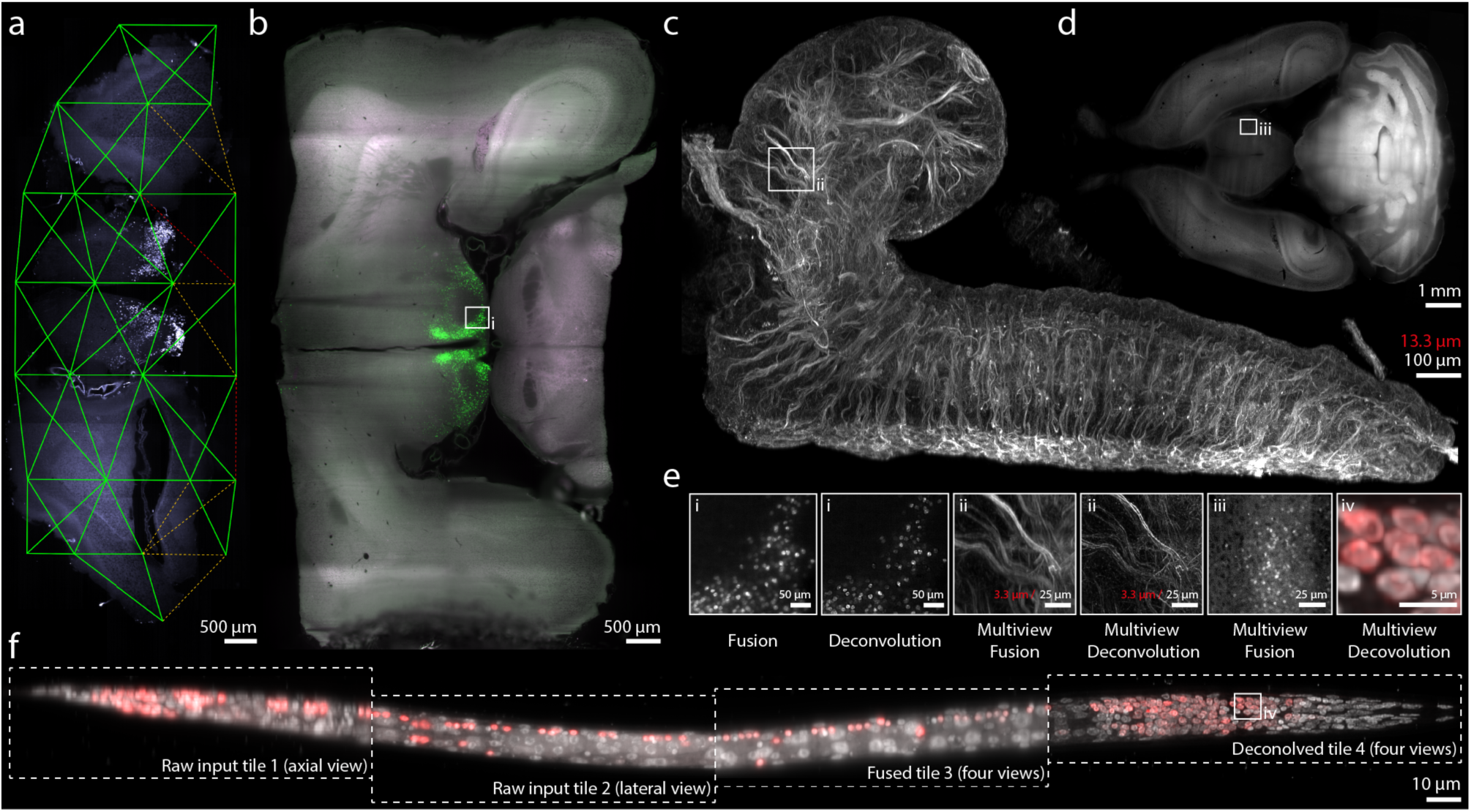
Reconstructed samples. **(a)** One slice through an acquisition of an adult mouse Bsx^H2BGFP^ coronal slice encompassing the hypothalamus. Green lines indicate strong links between overlapping image tiles, dotted orange lines refer to links rejected because of low correlation, and red lines illustrate links that were determined to be unconcise. **(b)** One slice through an adult mouse brain expressing an H2B-eGFP lineage tracing marker in BSX-expressing neurons, the box is highlighted in (e). **(c)** Maximum intensity projection of a central part of a 7.5-times expanded central nervous system of a *Drosophila* 1^st^ instar larva with immunostaining for tubulin (Alexa-488) and imaged with multi-tile IsoView light-sheet microscopy, boxes are magnified in (e). (**d**) One slice through a whole multi-tile, multi-view reconstructed adult Bsx^H2B-GFP/+^ mouse brain, the inset is highlighted in (e). **(e)** Zoom-ins to specific areas of (b,c,d,f) illustrating (sub-)cellular resolution and the advantage of (multi-view) deconvolution over (multi-view) fusion. **(f)** Fixed *C. elegans* dauer acquired in four tiles with four views each expressing tagRFP in all neuron nuclei, co-stained with DAPI. Boxes highlight the quality of axial and lateral raw input data, multi-view fusion, and multi-view deconvolution.

Acquisitions often consist of hundreds of image tiles, each many gigabytes in size showing very different information content (**Fig. 2**). We therefore compute pairwise overlaps using the parameter-free Phase Correlation (PC) method [ 12]. It computes all possible shifts between two images where intensity peaks in the resulting PC matrix correspond to shifts with high correlation (**Supplementary Note 3** and **Supplementary Fig. 6**). To accommodate large images, we can compute the PC matrix on downsampled images and localize peaks with subpixel precision [13]. Using simulations, we show that errors below 1 pixel can be achieved while reducing computation times 100-fold (**Supplementary Fig. 7-10**). All pairwise shifts (*links*) can be filtered by minimum correlation and distance from metadata defined positions and can optionally be interactively verified (**Supplementary Fig. 11**).

To compute final image tile locations, we developed a new optimization algorithm. It is based on identifying tile positions that minimize the distance between all links [14, 15]. Compared to computing a minimum spanning tree [16], normally distributed link errors (**Supplementary Fig. 10**) are averaged out during optimization since each tile is typically linked to many neighbors (**Fig. 2a**). To remove incorrectly computed links, we iteratively remove the link that disagrees most with the global optimization result using a new compound metric. So far, unconnected tiles (e.g. empty images) and multiple independent objects in an acquisition were handled by simply dropping them [14, 17], or assuming regular, 2d translational grids [18]. We developed a generic solution by introducing the concept of *strong* and *weak* links (**Supplementary Fig. 5**) that is independent of the tile arrangement and the transformation models used. Strong links are defined by confirmed image overlaps, while weak links are derived from approximately known tile positions (e.g. metadata). We first identify groups of tiles connected by strong links and compute their positions relative to each other for each group. Transformations within these groups are then fixed and final positions for all tiles are computed by minimizing the distance between all weak links (**Fig. 2a** and **Supplementary Fig. 5** and **Supplementary Note 4**).

To compensate for sample-induced light refraction, wavelength-dependent aberrations, and remaining small alignment errors we implemented an easy-to-use interest point based alignment step supporting affine transformations. We automatically extract interest points and apply a variation of the iterative closest point algorithm (ICP) [19] combined with our new global optimization algorithm. We thereby compensate smaller rigid or affine distortions including major effects of chromatic aberration if autofluorescence levels are sufficiently high (**Fig. 1i,j** and **Supplementary Fig. 12**).

Since emitted light is distorted by the sample, maximum imaging depth is limited. To overcome this problem, we acquire samples from opposing directions by rotation (**Fig. 1c**) or by simultaneous acquisition with two objectives [20]. We implemented a new algorithm to register large multi-tile *views*, where each *view* consists of a set of aligned image tiles from one physical orientation. We segment interest points in virtually fused, downsampled images of each multi-tile view and identify corresponding interest points using adaptations of geometric hashing [21]. This significantly improves matching performance and robustly aligns large volumes, effectively doubling the imaging depth of any sample (**Fig. 1c** and **Tab. 1**).

**Table 1:** Performance comparison. Comparison of BigSticher features and performance with other available stitching programs. Benchmarks were performed on a HP Z840 workstation running Windows 10 Pro with two Intel Xenon E5-2667v3 CPUs and 512 GB memory. The benchmarks for the Arivis software were performed on a HIVE system running Windows Server 2012 R2 with two Intel Xenon E5-2640v3 CPUs and 256 GB memory. The latest stable version of each stitching program was used. BigStitcher datasets were stitched using 4x times (x,y) downsampling, and fusion was performed at stated downsampling levels and saved as TIFF stacks. Correctness of the stitching could only be confirmed in the BigStitcher in full detail due to the flexibility of interactive inspection. Processing of multi-view, dual-illumination datasets as well as ICP refinement and virtual fusion is only possible in BigStitcher. Note that fusion in BigStitcher supports intensity adjustment and scales to arbitrary sizes. All displayed values are averaged from three independent runs of each respective software except those marked with a star. **#**Note that image fusion results are not comparable since all applications except BigStitcher fuse translation-only datasets, which is a significantly simpler and easy to optimize task.

| **Table 1: Performance comparison** |  |  |  |  |  |  |
| --- | --- | --- | --- | --- | --- | --- |
| Data Size | 130 Mb / 63 Gb / 300 Gb | 130 Mb / 63 Gb / 300 Gb | 130 Mb / 63 Gb / 300 Gb | 130 Mb / 63 Gb / 300 Gb | 130 Mb / 63 Gb / 300 Gb | 1.67 Tb |
| Software | Illumination Selection | Stitching<br>(depends on Illumination Selection) | ICP refine | Fusion# @ 8x down-sampling / full res.<br>(depends on Illumination Selection) | Virtual Fusion @ full res. | Multi-view Registration |
| BigStitcher | n.a. / 9 s / 14 s | 4 s / 1 min 28 s / 7 min 15 s | 9 s / 57 s / 2 min 52 s | 1 s / 9 s / 44 s<br>7 s / 1 h 23 min / 8 h 47 min | 1 s / 15 s / 19 s | 6 min 30 s |
| TeraStitcher | X | 16 s / 34 min 43 s / 2 h 29 min | X | 1 min 15 s / 14 min 23 s / 1 h 4<br>2 min 29 s / 26 min 41 s / 2 h 35 min | X | X |
| ImageJ Stitcher | X | 10 s / 61 min / 5 h 35 min | X | X / X / X<br>16 s / 10 h 36 min / 44 h 36 min | X | X |
| Xuv Tools | X | 2 s / X / X | X | X / X / X<br>9 s / X / X | X | X |
| Arivis | X | 31 s / 47 h 15 min / 7 d 4 h* | X | X / X / X<br>2 s / 5 min / 51 min | X | X |
| **BigStitcher benchmark for processing a terabyte-sized multi-view dataset** |  |  |  |  |  |  |
| Data Size | 1.67 Tb |  |  |  |  |  |
| Software | Illumination Selection | Stitching | ICP refine | Fusion @ 4x downsampling | Virtual Fusion @ full res.<br>(display / save) | Multi-view Registration |
| BigStitcher | 59 s | 1h 14 min | 7 min 28 s | 1 h 1 min | 50 s / 23 h 50 min* | 6 min 30 s |

For downstream analysis datasets can be fused or directly analyzed using BigDataViewer plugins [22]. We implemented an algorithm for real-time fusion by multithreaded processing of the currently visible plane in virtual images using blockwise multi-resolution loading, which can optionally be performed downsampled and on regions of the sample (**Supplementary Fig. 13**). It enables fusion of terabyte-sized images on machines with little memory (**Supplementary Fig. 14**), while increased memory and compute power leads to very fast processing (**Tab. 1**).

Deconvolution is an established method to increase contrast and resolution in light microscopy acquisitions [23,24] and required point spread functions (PSF) are typically estimated using fluorescent beads embedded with the sample [25]. To handle multi-tile, multi-view acquisitions, we extended deconvolution code [26] allowing BigStitcher to deconvolve selected regions and significantly improve image quality (**Fig. 2b-e**). To be able to apply this approach to cleared samples we developed a protocol for embedding fluorescent beads in polymerization solution enabling robust PSF measurement (**Fig. 2b,e**). We furthermore combine ExM with IsoView light-sheet microscopy [20] allowing acquisition of multi-view, multi-tile datasets of expanded tissues enabling reconstruction of entire *Drosophila* larval nervous systems with spatially isotropic subcellular resolution (**Fig. 2c,e**).

BigStitcher is a powerful software package that enables efficient and automatic processing of terabyte-sized datasets. It addresses major unsolved issues such as easy import, managing of large images, datasets acquired in a non-regular grid, globally optimal alignment of sparse datasets, illumination selection, multi-view alignment of multi-tile acquisitions, PSF extraction, and interactive fusion. The aligned dataset and all intermediate steps are interactively displayed. The user has the option to verify and interact with the alignment process at any time to confirm and potentially guide proper alignment of complicated datasets (**Supplementary Fig. 3,11,15,16**). Automatic reconstruction of even large datasets can be achieved within tens of minutes and BigStitcher clearly outperforms existing software in terms of performance, functionality, and user-interaction (**Tab. 1**) [14,16,17]. BigStitcher supports cleared samples (**Fig. 2a,b)**, ExM samples (**Fig. 2c,e** and **Supplementary Fig. 17**), standard 2D and 3D confocal and widefield acquisitions, as well as tiled, multi-view light-sheet acquisitions (**Fig. 2f**). BigStitcher is implemented in ImgLib2 [11], open-source and provided as a Fiji [27] plugin with comprehensive documentation <u>(http://imagej.net/BigStitcher).</u> It is compatible with the ImageJ Macro language for most of its functionality and can thus easily be automated. These properties make BigStitcher a powerful and scalable tool for the handling and reconstruction of tiled, high resolution image datasets acquired by new light microscopy technologies.

## Acknowledgments

We thank T. Pietzsch and S. Saalfeld for insightful discussions and BigDataViewer & ImgLib2 support, C. Rueden for Fiji support and maintenance, the Caenorhabditis Genetics Center at the University of Minnesota for providing C. elegans strains. S.P., F.R.R., F.P., M.T. were funded by MDC Berlin, D.H., H.H., H.L. were funded by LMU Munich and the Nanosystems Initiative Munich (NIM), P.T, N.R., R.K.C., A.C., P.J.K. were funded by HHMI Janelia.

## Author contributions

S.P. conceived the idea in discussions with H.H., H.L, and M.T. D.H. and S.P. developed the algorithms and implemented the software. F.R.R. performed all clearing experiments, reconstructions and benchmarks. F.P. imaged and reconstructed the C. elegans. P.T. and N.R performed ExM sample preparation. R.K.C. and P.J.K. developed the ExM-optimized IsoView microscope and imaged the sample. S.P. reconstructed the ExM sample. S.P., M.T., H.L, H.H., P.J.K, and A.C. supported and supervised the project. S.P., D.H., and F.R.R. wrote the manuscript with input from the coauthors.

## Methods

### Animals

For clearing we used a previously generated Bsx^H2BeGFP^ mouse line [28], where the exon 1 of the bsx gene is replaced starting at the ATG with the coding sequence for histone2B eGFP. Brains from 10-week old female Bsx^H2BeGFP/+^ mice were used for tissue clearing and imaging. *C. elegans* dauer larvae expressing tagRFP fused to a nuclear localizing sequence under the pan-neuronal rab-3 promotor in all neuron nuclei [29] were obtained by selecting dauer larvae in 1% SDS for 30 minutes [30]. Dauer larvae were fixed with 4% PFA for 30 minutes on ice, placed in 70% Ethanol overnight at 4°C and subsequently stained with DAPI. Experiments were conducted according to institutional guidelines of the Max Delbrück Center for Molecular Medicine in the Helmholtz Association after approval from the Berlin State Office for Health and Social Affairs (LAGeSo, Landesamt für Gesundheit und Soziales, Berlin, Germany). *Drosophila* larva used for ExM were obtained from the strain w;;attp2, carrying an empty attp2 landing site [31].

### Clearing and Expansion

Tissue clearing was performed using the CLARITY protocol [1]. Mice were deeply anesthetized by intraperitoneal injection of 100 mg/kg Ketamine and 15 mg/kg Xylazine. Mice were exsanguinated by transcardial perfusion with 25 ml cold PBS followed by whole body perfusion with 25 ml cold monomer solution (4 % v/v acrylamide, 4 % w/v Paraformaldehyde (PFA), 0.25 % w/v VA-044 in PBS). The brains were collected and fixed in monomer solution for 2 more days. Next, the whole brains were placed in fresh monomer solution and oxygen was removed from the tubes by vacuum and flushing the tube with nitrogen gas for 15 minutes. The samples were then polymerized by placing the tubes in a 37 °C water bath under gentle shaking for 2 hours. Polymerized brains were placed in clearing solution (4% SDS in 200 mM Boric acid). Brains were incubated in clearing solution for 1 week at 37 °C with daily solution change. Then, the brains were actively cleared using the X-Clarity setup from Logos Bioscience for 24 hours with a current of 1 A at 37 °C. Cleared brains were washed twice overnight with 0.1 % v/v Triton X-100 in PBS and once with PBS. Before imaging, brains were placed overnight in FocusClear for refractive index matching.

For ExM, the nervous system of a 1^st^ instar *Drosophila* larva of was extracted, fixed in 4% PFA/1xPBS/0.1%Triton for 1 hour and washed 2×10 min in 1xPBS/0.1% Triton. Before each antibody usage, the nervous system and the antibodies were blocked in 5% goat serum/1xPBS/0.1%Triton for one hour. Following the blocking, the nervous system was incubated overnight at 4°C in 1:500 monoclonal Anti-a-Tubulin antibody produced in mouse (Sigma Aldrich T6199 1mg/ml). After 5×10 min washing (1xPBS/0.1%Triton), the secondary antibody 1:250 Anti-Mouse CF™488A antibody produced in goat (Sigma Aldrich AB4600387 2mg/ml) was added overnight at 4°C. The stained brain was washed in 1x PBS and then processed using an Expansion Microscopy (ExM) method in which the gel recipe and procedure was modified to achieve 7,5-fold expansion in each dimension (unpublished, BioArxiv in preparation). Briefly, the specimen was treated with acryloyl-X as in standard ExM and embedded using a gel recipe modified from the original method [3]. The modified recipe uses a reduced cross-linker concentration to achieve greater expansion. After digestion with proteinase K as in the original method, a re-embedding step toughens up the gel, which would otherwise have poor mechanical properties.

### Imaging

3D images of cleared mouse brains were imaged using the Zeiss Lightsheet Z. 1 microscope. Each sample was attached to the sample holder using a cyanoacrylate-based glue. The mounted sample was placed in the FocusClear pre-filled imaging chamber. Images were acquired using the EC Plan-NEOFLUAR 5×/NA 0. 16 objective together with the LSFM 5x/NA 0.1 illumination objectives on a Zeiss Light-sheet Z. 1. The data was acquired using dual side illumination and from different angles. Images were collected with two 1920 × 1920 pixels sCMOS cameras and stored in the Zeiss CZI file format. Fixed *C. elegans* dauer larvae were embedded in 1.2% agarose containing fluorescent beads and imaged using the same microscope in a water-filled sample chamber. Imaging was performed using the 20x/ NA 1.0 objective with additional 2x zoom. 3D images from a cleared and expanded central nervous system of a *Drosophila* 1^st^ instar larva were acquired using an IsoView light-sheet microscope [20] that has been modified for multi-tile acquisition. To prepare the sample for imaging, excess gel surrounding the expanded sample was removed using a scalpel, leaving four flat and smooth gel surfaces for imaging. Some extra gel was left underneath the sample for mounting, and the sample was affixed to a cylindrical post using a cyanoacrylate-based glue. Mounted sample was placed in the imaging chamber filled with deionized water. Orthogonal views for each tile of the sample were acquired sequentially by switching the illumination and detection orders in IsoView. Images were acquired using SpecialOptics 16x/NA 0.71 objectives and recorded using full frames (2048 × 2048 pixels, pixel pitch of 0.4125 μm in sample space) of Orca Flash 4.0 v2 sCMOS cameras. The sample was held stationary during multi-view acquisition of each tile, and depth-sectioned images were acquired every 0.8125 μm by translating the detection piezos over a range of 750 μm. A tile for each view thus covered a field of 832 μm (X), 832 μm (Y), and 750 μm (Z). Automated tiling across the entire sample was achieved by moving the sample in predefined steps of 600 μm in X, Y, and Z until full coverage was achieved. Bi-directional light-sheet illumination was achieved using opposing SpecialOptics objectives and the illumination NA was chosen to be 0.0315 for a confocal parameter of approximately 416 μm. The light-sheets from opposing arms were shifted approximately by their Rayleigh length (208 μm) toward the illumination objectives so that each light-sheet provided uniform coverage of the respective half in the acquired image.

### PSF measurement and PSF Extraction

In light-sheet microscopy measured PSFs often differ significantly from simulated ones due to variable precision of light-sheet alignment in every experiment. Therefore, light-sheet deconvolution usually relies on the extraction of PSFs from the actual experiment [25, 26]. To be able to perform PSF extraction in cleared tissue we developed a new protocol. Estapor Fluorescent Microspheres (F-XC 030) were diluted 1:20000 in monomer solution containing bis-acrylamide (0,05 % v/v bis-acrylamide, 4 % v/v acrylamide, 4 % w/v Paraformaldehyde (PFA), 0.25 % w/v VA-044 in PBS). The monomer solution was polymerized under constant vacuum and shaking at 37 °C for 2 hours. The formed hydrogel was incubated in FocusClear overnight and imaged using the Zeiss Lightsheet Z. 1 microscope with the same experimental settings used to acquire previous samples. For *C. elegans* dauer imaging fixed larvae were embedded in 1.2% agarose together with Estapor Fluorescent Microspheres (F-Z 030), diluted 1:2000. For ExM data acquired on the IsoView microscope depth-sectioned images (0.4125 μm step) of fluorescent beads (200nm diameter) embedded in 0.6% low-melting-temperature agarose were imaged using the same experimental settings as for sample imaging. For all samples, PSFs were extracted by detecting interest points in the acquired bead images. Potential bead aggregates were excluded by manual removal on the maximum intensity projection using the BigStitcher module “Manage Interest Points > Remove Interactively”.

### Data processing pipeline

All data shown in this paper was processed using the BigStitcher Fiji plugin. Zeiss CZI files and TIFF files exported by custom microscopes were imported using the AutoLoader and subsequently converted to the HDF5 format. For Zeiss CZI files, approximate tile positions and rotation angles were imported automatically, for other files they were specified by hand using BigStitcher tools (**Supplementary Fig. 3, 16**). For each tile, the best illumination was selected. Tiles were aligned using the phase correlation method together with two-round global optimization, followed by ICP refinement. Interest point detection for each multi-tile view was performed. The fast descriptor-based rotation-invariant algorithm or the descriptor-based translation-invariant algorithm after manual rotation application were used to register the interest points of each angle, followed by another round of ICP refinement of all image tiles of the acquisition. Fused and deconvolved images were exported as TIFF files.

### Illumination selection

When imaging large samples using sequential dual-sided illumination, typically only illumination from one direction provides good image quality (**Supplementary Fig. 2**). We therefore implemented a simple *illumination selection* functionality in BigStitcher. It starts by *combining* all (selected) images by their illumination attribute, i.e. it groups images that share all other attributes besides illumination direction. In each of the resulting groups we select the best image. This is achieved by loading the pixel data for all images in the group at the lowest resolution level (in the case of non-multiresolution images, this corresponds to the original image) and calculating a *quality metric*. We currently offer mean intensity and mean gradient magnitude as quality metrics, which are typically sufficient for robust estimation of the higher quality illumination direction (**Fig. 1d**). The image with the highest score is kept, while all other images are marked as *missing* in the dataset, which will lead to them being ignored in subsequent processing steps. Optional resaving of the dataset after this step potentially decreases storage requirement two-fold. Prior to applying automatic illumination estimation, the user has the option to verify and potentially change the result.

### Pairwise Stitching using Fourier-based Phase Correlation

We calculate pairwise translational shifts using our ImgLib2 [11] implementation of the Fourier-based *phase correlation* algorithm [12]. In noiseless images, the method produces a phase correlation matrix (PCM) *Q* containing a single δ-impulse at the location corresponding to the shift between the two images. Real images might contain multiple peaks (**Supplementary Fig. 6**) and we localize the *n* highest peaks in *Q* by detecting peaks with subpixel accuracy using a *n*-dimensional implementation of a quadratic fit [13]. Aside from allowing subpixel-accurate registration, we can use the precision obtained from the subpixel accuracy of the phase correlation to counteract the effects of downsampling, allowing us to achieve registration of similar quality to full-resolution with significant performance gains (**Supplementary Fig. 8-10**). Due to the periodic nature of the Fourier shift theorem, each peak in the PCM actually correspond to 2^n^ possible shifts in *n* dimensions. We therefore test each of these candidate shifts by calculating the cross-correlation between the images *I_1_* and *I_2_*, optionally with interpolation in the case of sub-pixel shifts. We choose the shift vector *t* corresponding to the highest cross correlation as the final result after applying downsampling correction, if necessary.

It is often necessary to not only align two single images but groups of images, e.g. all channels of a tile. We therefore implemented a flexible framework for the registration of grouped images (**Supplementary Note 3**). The two images *I_1_* and *I_2_* can have arbitrary affine pre-registrations such as sample rotation, correction of axial scaling or already performed registration steps. If pre-registrations of *I_1_* and *I_2_* are identical or are only based on different translations or axis-aligned scalings we run the phase correlation on (downsampled) raw input images, otherwise on virtually fused images (**Supplementary Note 3**).

### Downsampling and Simulations

To assess the effect of downsampling on the pairwise stitching we use simulations of spheroid-like objects at different signal-to-noise ratios (SNRs) as ground truth. We create realistic images by mimicking image creation in light-sheet microscopy including optical sectioning, 3-fold anisotropy, light attenuation, convolution, and pixel intensity generation using Poisson processes [26]. Importantly, pairs of overlapping images that we use for benchmarking the subpixel phase correlation method are created using different Poisson processes and are additionally rendered with half a pixel offset of the full resolution images to avoid nearly identical overlaps at high SNRs due to the simulation process (**Supplementary Fig. 7**). We simulate 500 pairwise overlaps, each at SNRs ranging from 1 to 32, and lateral downsamplings ranging from 1x to 8x where axial downsampling is matched as good as possible to achieve near-isotropic resolution as in the actual software. We illustrate that across SNRs downsampled images yield a constant registration quality, which even exceeds that of registration at full resolution for low SNRs. This is achieved through a combination of the smoothing effect during downsampling (**Supplementary Fig. 7**) and precise subpixel-localization (**Supplementary Fig. 8-10**). Registrations with a constant quality of an average error of below one pixel can be computed at a fraction of the computing time compared to full resolution, typically 4 - 120 times faster. Existing outliers are filtered during global optimization and overall registration quality can further be improved during the ICP refinement step.

### Global optimization

To calculate the final transformations of each image tile we extend the concept of globally optimal registration by iterative minimization of square displacement of point correspondences (**Supplementary Note 4**) [14,15]. We express pairwise shifts as point correspondences between the bounding box vertices of the overlap region of the images and the same points transformed by the inverse pairwise shift. We globally optimize the registrations *R* in all connected components *CC* of the link graph of images *V* and (strong) links *C*, given point matches (corresponding points) *PM* and fixed images *F* by minimizing:

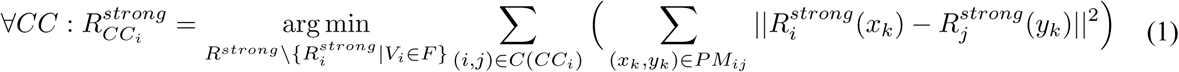

In some cases, erroneous pairwise links might not have been filtered out, e.g. due to medium cross-correlation, repetitive patterns, or a low number of correspondences in the ICP refinement. This leads to persistently high registration errors after global optimization, which manifests in a large distance error, i.e. the difference between the individually computed distance between of images (link) and the actual distance between them after global optimization. Iterative removal of the link with the highest distance error from the link graph and repeating the global optimization leads to convergence to user-defined thresholds [14]. We extend this concept to affine transformations, introduce a new heuristic that additionally incorporates link quality and implement it in an extendable framework required for the two-round global optimization (**Supplementary Note 4**).

If the dataset contains empty tiles or multiple disconnected objects with image tiles that do not have links between them, the final transformations will not be propagated between them (**Supplementary Fig. 5**). We therefore developed a two-round global optimization that is capable of aligning independent connected components of the link graph using weak links defined by pre-existing transformations (e.g. approximate locations from metadata or manual alignments). For that purpose, we use the corners of the bounding box of their overlap region, transformed using the results R_strong_ of the first round (eq. 1), as point correspondences. The between-component transformations can then be determined by minimizing:

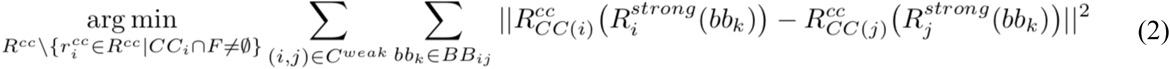

The final transformations are given by concatenating the in-component and between-component registrations. Using the two-round strategy, registrations are propagated between connected components and distances between neighboring objects are preserved as-well-as-possible (**Supplementary Note 6** and **Supplementary Fig. 5**).

Our global optimization is agnostic to the nature of the point correspondences and transformation model, which allows us to use the same algorithm for translation-based alignment of for example tiled datasets using phase correlation, as well as affine registrations of multi-tile multi-view datasets based on ICP refinement or geometric descriptor matching.

### Iterative Closest Point Refinement

Although the Phase Correlation-based image stitching produces relatively high-quality alignments, smaller errors can remain due to its inaccuracy (**Supplementary Fig. 8-10**). Furthermore, it is not able to correct for non-translational effects such as chromatic aberration or sample-induced light refraction. These effects can be better approximated using affine transformations. We therefore automatically detect interest points and run an Iterative Closest Point algorithm [19] for each overlapping pair of images, where the assignment of correspondences is limited by a distance threshold. We use the identified corresponding points of all pairwise links and compute a globally optimal affine transformation for each tile using our new global optimization algorithm. To avoid scaling of datasets, we regularize the affine transformation using a rigid transformation [ 15]. The resulting alignment usually improves the alignment quality and the same strategy can be applied to multichannel alignment if sufficient autofluorescent signal is available (**Supplementary Fig. 12**).

### Geometric Local Descriptor Matching

To identify corresponding interest points in between two point clouds, geometric local descriptor matching has been proven to be a powerful technique [21, 32]. The basic idea to express each interest point as a geometric constellation using its *n* (typically three) nearest neighboring interest points. The vector difference between two descriptors then describes how similar the local area of two points is. A geometric local descriptor (GLD) is assumed to be a correspondence candidate if it is at least *m* (typically one to ten) times more similar than the second most similar GLD [13]. True corresponding interest points between two point clouds are finally identified using the random sample consensus algorithm [33] on a regularized affine transformation model. However, fast GLD matching using the rotation-invariant technique based on geometric hashing [21] requires relatively randomly distributed points to robustly identify correspondences, while the non-accelerated, redundant, translation-invariant counterpart [32] identifies correspondences reliably in non-rotated point clouds of only up to a few thousand points in reasonable time. Here, we extended both techniques to better suit the requirements when attempting to identify corresponding interest point in between point clouds of prior unknow size derived from imaged structures that are potentially rotated relative to each other.

Redundancy is a powerful mechanism for GLD matching. It uses additional nearest neighbors but excludes some of them sequentially during matching making it more robust to potentially mis-detected interest points [32]. We therefore extend the fast rotation-invariant technique based on geometric hashing [21] with the capability for redundancy. This significantly increases the chance of being able to align randomly oriented point clouds very fast, albeit at low inlier ratios (ratio of true correspondences to total number of correspondence candidates).

Rotation invariance is not desired if both point clouds are known to be approximately in same orientation, for example if the rotation of the sample performed by the microscope was known and has been applied to the dataset. Checking for potential rotations simply increases the chance for wrong correspondence candidates. We therefore implemented a fast translation-invariant GLD based on geometric hashing that supports redundancy. All four versions of GLD are available in BigStitcher to enable robust multi-view alignment.

### Virtual Image Fusion

A set of overlapping, transformed image tiles are fused into one output image using a per-pixel weighted average that minimizes boundary artefacts and can increase contrast by incorporating entropy estimation (**Supplementary Note 6**) [21]. To correct for unequal brightness and contrast in adjacent images, we optionally perform adjustment of the pixel intensities using a linear transformation per image. An optimal adjustment can be estimated using the same optimization framework used for image registration (**Supplementary Note 7**) [34]. The memory requirements for the fusion of large volumes can easily exceed the available RAM on a machine due to the size of the output and the combined size of the input images. We therefore developed a framework based on ImgLib2 *RandomAcessibleIntervals* [11], intensity transformations and coordinate transformations that virtually fuses all pixels of a defined bounding box using all input images and their associated weights. Since the input images are provided through virtual image loading, the size of a virtually fused image is close to zero, irrespective of the size of input and output images. Ideally, input images are available in blocks so that affine transformations that slice input images in arbitrary orientations do not require to load the entire image [9]. The output image can now be rendered on a pixel-by-pixel basis with minimal memory requirements. Additional caching of the input image and the output images allows an efficient multithreaded fusion for as-fast-as-possible processing given the available memory. Therefore, more RAM will effectively speed up the fusion process (**Tab. 1**), but even machines with very low RAM will be able to fuse terabyte-sized volumes (**Supplementary Fig. 14**). Fused images can be saved by choosing cached or virtual fusion and subsequently saving the ImageJ virtual stack using “Save as image sequence…”. Downsampling of the output can easily be incorporated by scaling the bounding box and pre-concatenation of the downsampling transformation with each image transformation. If the input files are multi-resolution, we automatically compute the optimal resolution level at which the input needs to be loaded. To optionally further reduce the image size of the fused image, the GUI offers to conserve the original anisotropy between lateral and axial of the acquired sample, which is a sensible choice if the dataset contains a single or opposing (e.g. 0 and 180 degrees) multi-tile views.

### Deconvolution

In addition to real-time image fusion, we offer deconvolution of bounding-box-defined volumes using a multi-view formulation of the iterative Richardson-Lucy deconvolution algorithm [23,24] with Tikhonov regularization and various optimizations [26]. The PSFs required for deconvolution can be extracted from interest points detected in the images (e.g. when subdiffraction fluorescent beads were incorporated with the sample) or supplied as TIFF stacks with odd dimensions by the user. BigStitcher offers GPU acceleration of the deconvolution on CUDA-capable Nvidia GPUs.

To allow deconvolution of multi-tile views, we extended the original deconvolution [26] to be based on the *virtual fusion*. Thereby, any number of input image tiles are virtually fused and serve as one of input views for the multi-view deconvolution. Proper multi-view deconvolution of partly overlapping samples requires sophisticated weight normalization in between views [26], which we implemented to be computed virtually. Since also the input views are also virtually loaded, the memory requirement of the deconvolution solely depends on the output image size and shows a significantly increased memory-efficiency. All virtual inputs and weights are additionally cached, ensuring highest-possible processing performance for systems with large amounts of RAM.

### Macro automation and headless operation

In addition to the graphical user interface (GUI), we offer standalone Fiji plugins for most of the individual steps, such as data import, illumination selection, pairwise shift calculation, link filtering, multi-view alignment, global optimization and image fusion/deconvolution. In macro mode results will not be displayed interactively but are instead saved to the XML project file or output files immediately. The individual steps can be recorded as ImageJ *macros* and easily combined into a script for headless batch processing [35].

